# Scientific Figure Library connects visual examples with reusable plotting code

**DOI:** 10.64898/2026.11.09.765504

**Authors:** Jiaxuan Lai, Yu Qiao, Xi Chen, Shiming Xie, Zhougeng Xu, Shipeng Guo

## Abstract

Code-generating assistants can adapt plotting scripts, while researchers select visual examples and assess their relevance. Reusing those examples requires the image, implementation, and input context to remain connected. Scientific Figure Library (SFL) organizes visual examples, plotting code, input descriptions, and provenance as reusable figure assets. A user-selected local library retains working revisions and published releases, while a common provider interface makes local and externally maintained collections available through the same discovery workflow. Researchers browse candidate figures, inspect an exact preview, confirm a selection, and materialize the corresponding assets into a project for adaptation and execution. Local clients and external agents access a shared service, connecting visual selection with code-based analysis across research environments. An accompanying open resource, ScientificFigureLibrary-personal, supplies modular examples through the Open Figure Modules provider. Its organization combines previews, reusable scripts, example inputs, and licensing information, making it a concrete resource for learning and adapting scientific visualization patterns. We describe the asset model, retrieval and confirmation workflow, execution handoff, and resource organization, supported by implementation inspection and automated software checks. In a public lung single-cell case study, we reused a cell-composition template for 32 idiopathic pulmonary fibrosis and 28 control donors and verified the donor-level inputs and plotted proportions. SFL provides a practical connection between finding a figure and obtaining the materials needed to work with it, while preserving the identity and provenance of the selected example.

## 1 Introduction

Scientific figures condense measurements and analysis into a form that readers can inspect and interpret. Their effectiveness depends on the question, the audience, and the choice of visual encoding [1]. Reusing a well-designed example offers a starting point for a new analysis, provided that its code, input requirements, and transformations remain accessible. Preserving these connections is part of the wider practice of recording the materials and steps that produce computational results [2,3]. Tool-using language-model agents extend the ways in which researchers can work with those materials. ReAct interleaves reasoning with external actions and observations, while CodeAct uses executable code to compose actions and respond to execution feedback [4,5]. In biomedicine, Biomni connects planning and code execution with an environment of tools, databases, and protocols; BioAgents combines specialized agents informed by tool documentation with retrieval over workflow documentation to support analysis [6,7]. These systems make structured resources and executable implementations relevant inputs to an agent-assisted workflow.

Visualization research has developed corresponding interfaces for creating and revising figures. LIDA organizes visualization-code generation into a multi-stage pipeline, and Data Formulator 2 combines graphical and natural-language input with data transformations and iteration history [8,9]. WaitGPT and DynaVis respectively address how users inspect generated data operations and make further visual edits [10,11]. Together, these studies distinguish several design problems: generating a figure, revising its data and appearance, and keeping the process understandable to its user.

A complementary question concerns the example that enters such a workflow: how should its image, implementation, expected inputs, and provenance be kept together so that it can be selected and reused in another project? The relevant object is a reusable figure asset that persists beyond an individual analysis session. Its organization connects visual recognition with the identification and stewardship of research software [12,13].

Here we present Scientific Figure Library (SFL), a local-first library that connects scientific figure examples with their reusable code and context [14]. Its workflow combines visual browsing, exact selection, versioned publication, and project materialization through a shared service for local clients and external hosts. The accompanying Open Figure Modules collection supplies modular examples from the ScientificFigureLibrary-personal repository [15]. We describe the design, its relationship to agent-assisted visualization, the organization of the open resource, and the handoff from selected assets to project execution.

## 2 Related work

### 2.1 Generation and iterative authoring

Declarative visualization languages make chart design explicit. The layered grammar of graphics describes plots through composable components, while Vega-Lite extends a high-level grammar to interactive visualizations [16,17]. Such representations provide structured targets for both human programming and model-generated code, as illustrated by their use in Data Formulator 2 and DynaVis [9,11].

LIDA connects data summarization, goal exploration, and visualization-code generation, including code refinement and execution [8]. Data Formulator 2 combines graphical specification with natural-language instructions and AI-generated data transformations. Its data threads retain iteration history so that users can revisit branches and build on previous results [9]. These systems address the creation and evolution of a visualization during analysis. SFL concentrates on discovering and transferring an identifiable example together with its reusable materials.

### 2.2 Inspection and user control

Inspection can act on different parts of an AI-assisted workflow. WaitGPT converts generated analysis code into an interactive representation of data operations, supporting monitoring and modification as the analysis unfolds [10]. DynaVis combines natural-language editing with synthesized persistent widgets, allowing users to explore and repeat visualization edits through direct interaction [11]. These systems offer concrete interfaces for examining or steering computational work. SFL places its inspection point at resource selection: the researcher or an explicitly delegated host reviews a specific preview before the associated assets are transferred. The research environment then supports execution-time inspection and subsequent editing.

### 2.3 Persistent resources and cross-project reuse

Reusable computation also depends on the organization of its supporting resources. Computational notebooks bring code, results, and explanation together within an analysis [18]. The nf-core community organizes reusable analysis pipelines through shared templates, documentation, tests, and contribution practices [19]. Its focus on complete workflows differs from a figure module, but illustrates how common structure makes an open software collection usable by others.

Persistence is already part of interactive authoring: Data Formulator 2 retains data threads, and DynaVis retains editing widgets [9,11]. SFL organizes a different persistent object, linking a figure preview to code, input descriptions, provenance, and a published release. Its Library retains these objects independently of a project, and provider-qualified selection identifies the example entering a new analysis. Open Figure Modules provides concrete packages for this asset model. The resulting focus connects interactive visualization with FAIR principles, software identification, and research-software stewardship [12,13,20]. It complements the composition and execution of tools by an external host with an explicit asset-management and handoff layer.

## 3 Design and implementation

### 3.1 A figure asset as the unit of reuse

The basic object in SFL is a Figure Unit: one or more visual assets, a canonical preview, and descriptive metadata. A plotting template adds code assets, implementation information, and links between code and figures. A visual-reference entry can retain an image and its context before executable code is available. This representation accommodates both the collection of useful examples and the later development of reusable implementations.

Metadata describes what a figure is for, what inputs it expects, and how its visual structure is organized. These descriptions connect a biological or analytical question with a plotting pattern. The image supports visual recognition; the code exposes the implementation; provenance and licensing record where the material comes from and how it can be reused. Keeping these elements together applies research-software identification and stewardship principles at the scale of an individual figure asset [12,13,20].

SFL stores assets in a global Library directory explicitly selected by the user. The Library is shared across projects, while each project receives only the assets selected for that task. This organization separates an accumulated collection of reusable knowledge from the transient files of an individual analysis. The local directory contains the authoritative asset records; indexes and preview caches support access to those records.

### 3.2 Publication preserves the relationship between an image and its code

A template series groups successive revisions of an asset. Editing creates a new content revision, and publication creates an immutable release associated with a review snapshot. The series retains pointers to its current working and published states, while earlier releases remain available for reference. Researchers can therefore improve an example without changing the identity of an already selected release.

Review combines structural checks with decisions about the figure–code association, the canonical preview, and the available evidence. Blocking issues are resolved before publication, and non-blocking warnings remain attached to the published object. The resulting release carries the content and review context that were approved together. This is particularly useful when a figure has multiple implementations or when a later revision changes its inputs or appearance.

### 3.3 A shared service connects local interfaces and external agents

SFL separates its business operations from the interfaces through which users access them (Figure 1). A shared TypeScript and Node service handles asset storage, retrieval, preview, publication, and materialization. Local clients call this service through a local interface. External coding assistants and research environments access it through a Model Context Protocol (MCP) adapter, with an optional MCP App providing visual interaction in supporting hosts [21].

**Figure 1.**
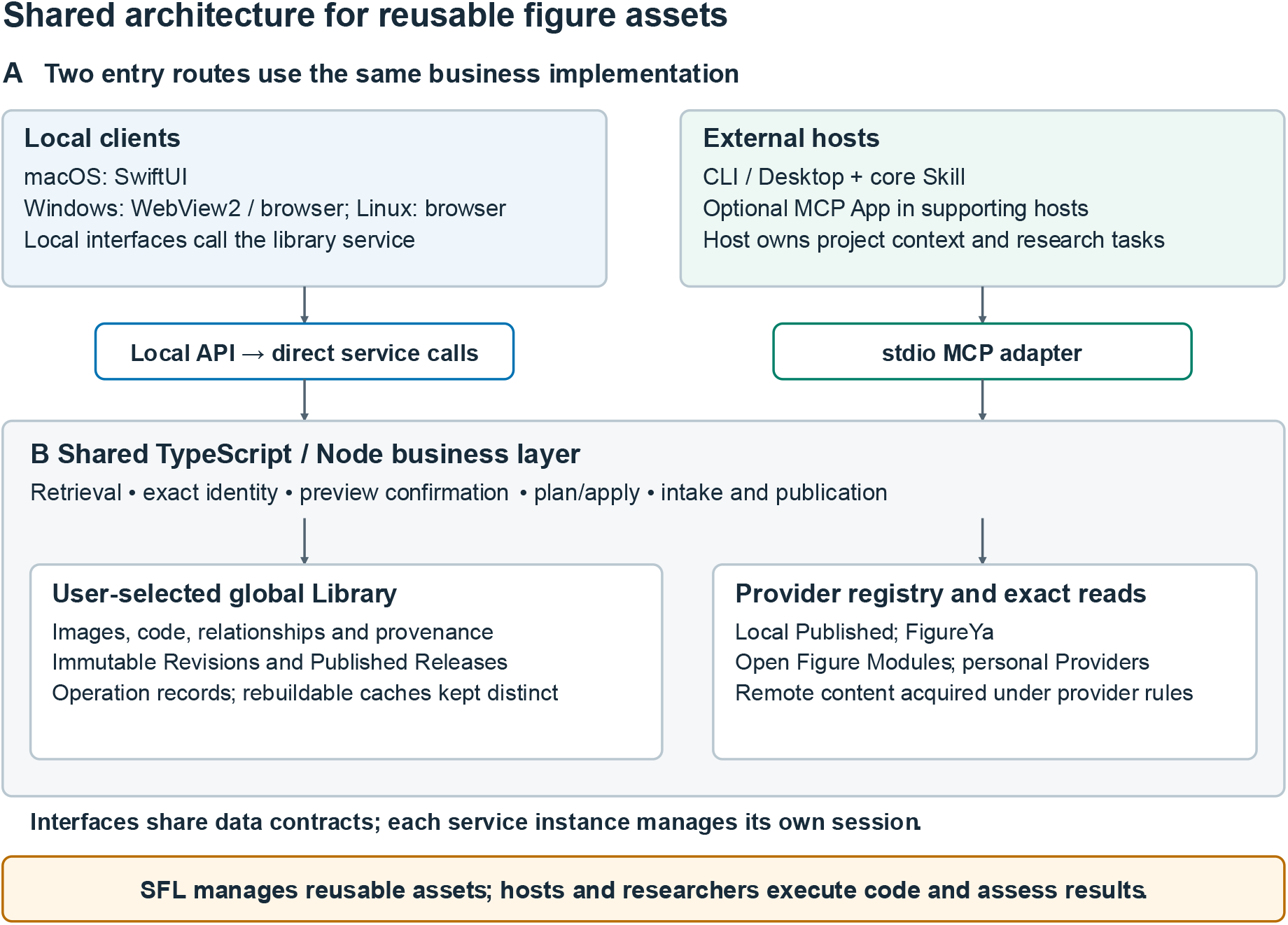
Shared architecture for reusable figure assets. (A) Local clients and external CLI/Desktop hosts access a common TypeScript/Node business implementation through separate interfaces. Local clients call the library service directly; external hosts use Model Context Protocol (MCP) and the core Skill, with an optional MCP App where supported. (B) The shared service handles retrieval, precise asset identity, preview confirmation and controlled writes. A user-selected global Library preserves images, code, their relationships, immutable Revisions and Published Releases. Providers connect the service to Local Published, FigureYa, Open Figure Modules and configured personal sources. Each service instance manages its own session. SFL organizes the reusable assets; the researcher and host retain project execution and interpretation.

The local-client architecture includes a native SwiftUI interface on macOS and Node-backed web interfaces on Windows and Linux. The same asset model and operations are available to external hosts through MCP. A core Skill explains the intended workflow to an agent, while the service enforces object identity and write conditions. This division allows interface-specific interaction without duplicating the rules that govern the Library.

The host supplies the reasoning and execution environment for a research task. It can interpret a request, inspect the selected code, adapt it to project data, and run it using an appropriate R or Python environment. SFL supplies the reference assets and records their transfer. Keeping these roles explicit makes the library useful to both manual workflows and agent-assisted analysis.

### 3.4 From candidate discovery to an exact selection

Discovery spans locally published assets and external providers. Local Published, FigureYa, Open Figure Modules, and explicitly opted-in personal providers retain distinct identities within the common search interface. Server-side lexical retrieval uses weighted descriptive fields, including titles, tags, visual and data profiles, and scientific-question metadata. A host can supplement this retrieval by inspecting the catalog and reasoning over candidate descriptions.

Search returns candidates for comparison. Exact selection then binds a candidate to its provider and the corresponding asset identifiers (Figure 2). For a local release, those identifiers include the template, revision, release, and content digest. For a module archive, the selector also captures the relevant source, archive, preview, and catalog identities. A selected figure is consequently identified by its content and source, rather than by its displayed title alone.

**Figure 2.**
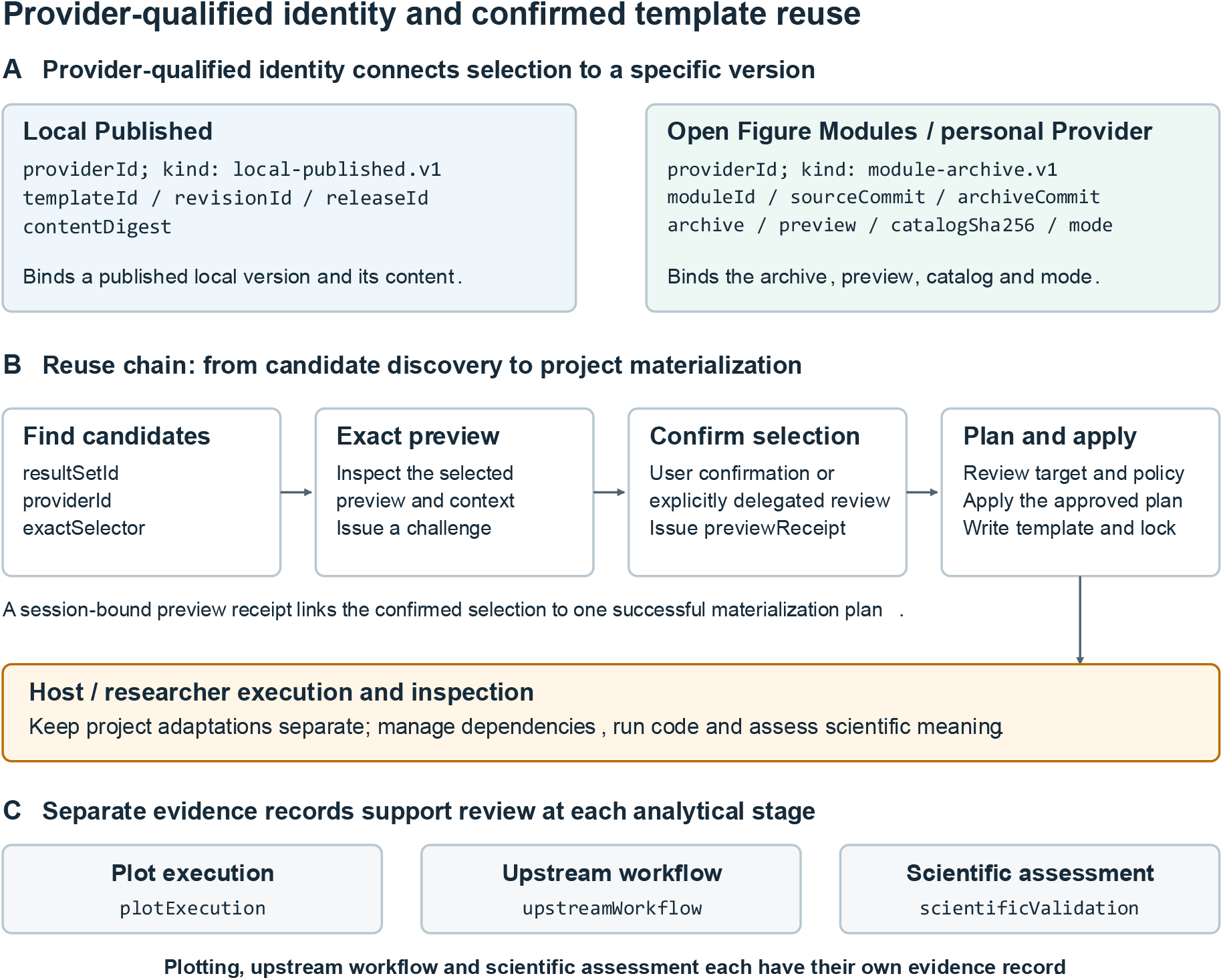
Provider-qualified identity and confirmed template reuse. (A) Example selector fields bind a Local Published object to its template, revision, release and content, or bind a module archive to its source/archive commits, preview, catalog and selected mode. The provider identifier supplies the source context. (B) Discovery proceeds through exact preview, confirmation and a reviewed materialization plan. The session-bound preview receipt is consumed by one successful plan; apply writes the selected template and lock metadata to the project. A supporting interface can obtain direct user confirmation, while the headless route follows user selection or explicit delegation. The researcher and host then adapt and execute code in the project environment. (C) Separate records for plot execution, upstream analysis and scientific assessment make the scope of supporting evidence available for review.

The researcher or an explicitly delegated host inspects the exact preview and confirms the choice. SFL issues a session-bound confirmation receipt that links this decision to subsequent materialization planning. Candidate browsing and exact confirmation thus serve different purposes: browsing explores the collection, while confirmation identifies the particular asset to be reused.

### 3.5 Materialization connects the Library to an executable project

Materialization writes the selected assets and a provenance lock into a specified project directory. Before writing, SFL presents a plan describing the asset, destination, and acquisition policy. Application checks the plan against the current Library and target state. Completed-operation records support checked replay and preserve the outcome of a write. Publication follows its own planned workflow, connecting a proposed asset, review decisions, and the resulting local release.

Once materialized, the files become working materials for the project. The host can inspect input tables, adjust aesthetic mappings, revise labels, or change the analysis while retaining a record of the starting template. Execution and analysis records then belong to the project, following the same principles of explicit inputs and computational history used in reproducible research [2,3,18]. This handoff is the point at which a stored example becomes part of a new analysis.

SFL records plotting execution, upstream-workflow evidence, and scientific assessment separately. These fields make the scope of available evidence visible when an asset is published or reused. Plotting-code execution remains with the host; a materialization receipt records the transferred assets, not the outcome of running them.

## 4 An open collection of reusable figure modules

### 4.1 The resource complements the client

ScientificFigureLibrary-personal is the public repository behind the Open Figure Modules content source [15]. Despite the repository name, it serves as a distributable collection of reusable scientific plotting examples. The SFL client manages discovery, selection, and local asset handling, whereas the resource repository maintains module content and its release catalog. This separation allows the collection to develop independently of the client. For a researcher, the preview and description expose the intended visual pattern; for an external host, the file inventory, input description, and code provide concrete materials to inspect before adaptation.

At inspection, the repository contained 182 module directories, matched by 182 archive-manifest entries. The collection is organized as individual modules rather than one monolithic script bundle. A module can bring together preview images, descriptive metadata, an organized plotting script, example input data, and provenance or licensing records (Figure 3). Catalog entries expose the information needed to find an example, while module archives carry the materials needed for its selected reuse.

**Figure 3.**
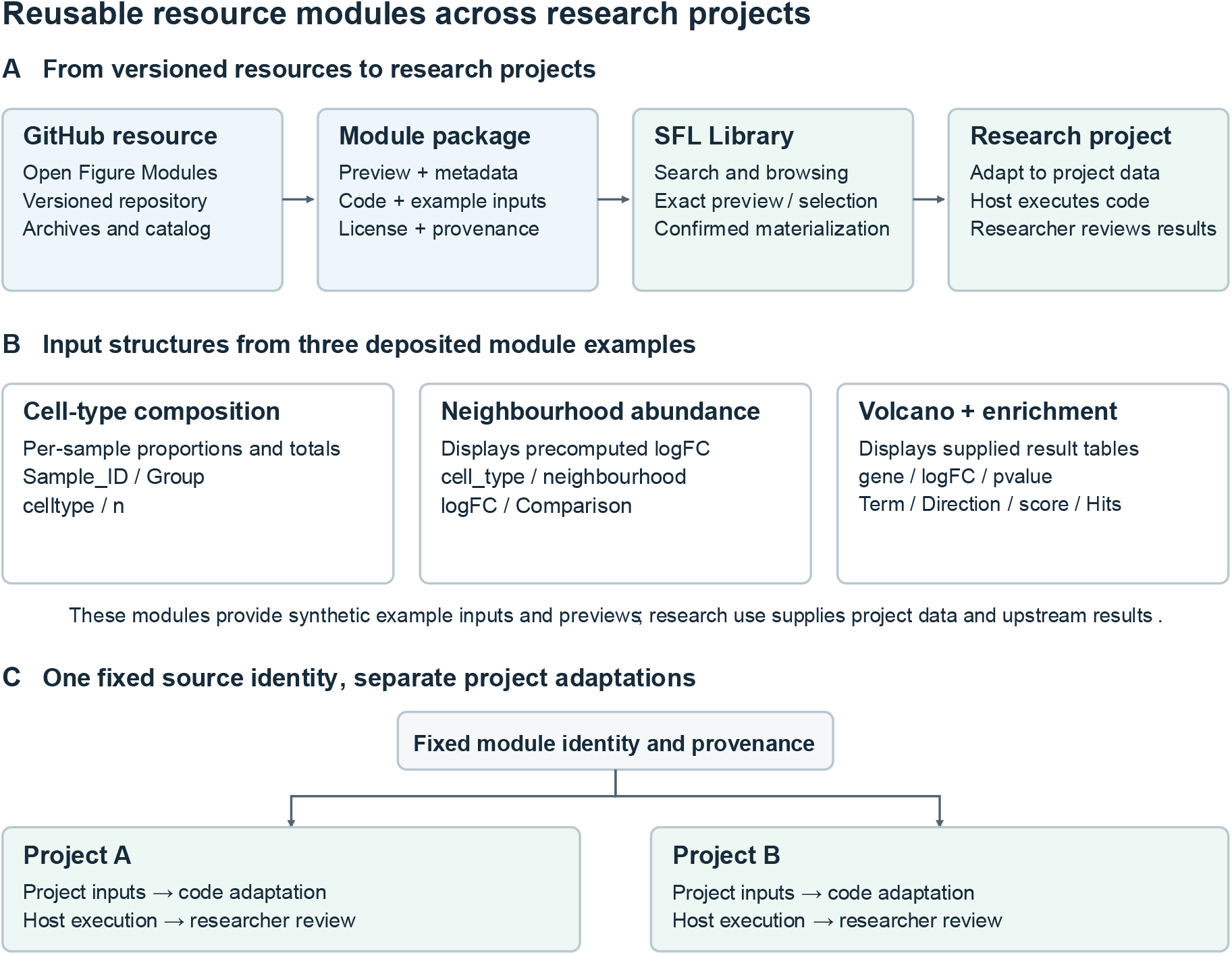
Reusable resource modules across research projects. (A) Open Figure Modules supplies versioned resources comprising previews, metadata, plotting code and, where supplied, example inputs, together with license and provenance information. SFL supports catalog discovery, exact selection and confirmed materialization; code adaptation and execution take place in the research project. (B) Three inspected modules expose distinct input structures: sample–cell-type counts for composition displays (`sc-celltype-sample-stacked-proportion`), precomputed neighbourhood effects for beeswarm displays `neighbourhood-abundance-beeswarm`), and gene-level and functional-summary tables for coordinated volcano/enrichment views (`volcano-inset-enrichment-bars`). Their distributed inputs are synthetic plotting examples, and each declares MIT code and CC BY 4.0 synthetic content/documentation. (C) The cross-project schematic shows how one fixed source identity can support separate project-specific adaptations while retaining provenance. Projects A and B illustrate this reuse model rather than denote evaluated biological datasets. Module source files and input schemas were inspected at the pinned resource snapshot; plotting scripts were not rerun for this figure.

### 4.2 Module structure makes an example inspectable

The module structure supports a sequence of practical questions. The preview shows the intended visual result; metadata describes the use case and data profile; input files show the expected shape of the data; and the plotting script exposes the transformations and graphical choices. Source and licensing records place the example in its distribution context. The combination helps a researcher move from recognizing a useful visual pattern to identifying what must change for a new dataset.

Three inspected modules illustrate different interfaces between analysis outputs and visual design [15]. The sample-level cell-composition module accepts a sample-by-cell-type count table and calculates within-sample proportions, displaying total counts and group facets. The neighbourhood-abundance module arranges precomputed neighbourhood log-fold changes by cell type and comparison. The volcano-and-enrichment module combines gene-level effect and significance values with direction-specific functional summaries. A researcher can inspect each schema, substitute a suitably prepared result table, and adapt the organized R script to the project. These modules package the presentation of analysis outputs rather than the upstream statistical procedures.

Provenance records distinguish external visual inspiration from the plotting-layer implementation distributed with a module. This makes the origin of a design and the scope of its reusable implementation inspectable. Synthetic example inputs serve as teaching and demonstration materials; they are identified as such in the module evidence rather than presented as observations from the source study.

### 4.3 Distribution links open resources to local work

Open Figure Modules uses a catalog and archive distribution path. SFL can discover entries from a bundled catalog and activate verified catalog updates through a signed feed. The complete archive is obtained for the selected module during an authorized acquisition or materialization step. The asset retains its provider-qualified identity through this process.

This arrangement gives the open collection and the local Library complementary roles. The open collection provides shared examples; the local Library retains the researcher’s reviewed assets; and a project receives the files needed for a particular analysis. An adapted result can later be imported and reviewed as a new local asset. Reuse therefore forms a cycle of discovery, adaptation, and contribution rather than ending with the download of a script.

## 5 Implementation checks and use in analysis

We inspected the storage model, provider selectors, preview-confirmation mechanism, materialization workflow, and associated automated tests. The test suite completed with 480 passing tests and three platform-dependent skips, and the build and MCP smoke checks completed successfully. The exercised paths included intake, immutable publication, exact-preview confirmation, materialization, and checked replay. Detailed execution records and source snapshots are retained separately from the conceptual description.

The inspected modules support the following reuse workflow. A researcher can first choose a visual pattern that matches the analytical question, then read its data description and code. After confirming the preview, the researcher can materialize the module, map project data onto its required inputs, and execute the adapted script in the host environment. The host and researcher retain the original template identity alongside the project’s modifications and outputs. This sequence connects the library’s visual interface with the code and data practices required for a reproducible analysis [2,3,18].

### 5.1 Public-data application: lung cell composition in IPF

We used the public IPF Cell Atlas (GSE136831) to examine donor-to-donor variation in annotated lung cell composition in idiopathic pulmonary fibrosis (IPF) and control lungs [22,23]. We reused the deposited cell annotations, cross-checked subject and disease labels, and excluded COPD and records the authors labeled Multiplet. We did not recluster cells, rerun doublet detection, or use the expression matrix. We aggregated retained cells by donor and cell type, completed zero-count donor–type combinations, and divided each count by the number of retained annotated cells in that donor. The denominator was all retained annotated cells, not just epithelial cells.

Using SFL in an isolated library, we retrieved the Open Figure Modules cell-composition template, inspected and confirmed its exact preview under the delegated selection, and materialized it to a case workspace. We preserved the original template and its identity record. In the host R environment, we mapped the donor counts to the template input and ran the adapted plotting code. SFL did not perform the statistical analysis. Independent aggregation checks reconciled the counts and the figure’s underlying proportions with the deposited annotations (Supplementary Material S1).

The analysis included 32 IPF donors (144,404 cells) and 28 control donors (95,303 cells), with 38 observed cell-type labels. Figure 4A shows the variation in composition among donors. For the prespecified descriptive AT2 comparison, the median fraction was 0.17% (IQR 0.05–0.36%) in IPF and 0.56% (IQR 0.28–1.26%) in controls (Figure 4B). The difference in medians (IPF minus control) was -0.39 percentage points (95% percentile bootstrap interval -1.07 to -0.12; 10,000 resamples of donors within groups). This was an unadjusted, exploratory contrast in the public cohort. Including the author-labeled multiplets in the denominator gave a difference of -0.39 percentage points. Neither comparison establishes a diagnostic or mechanistic finding.

**Figure 4.**
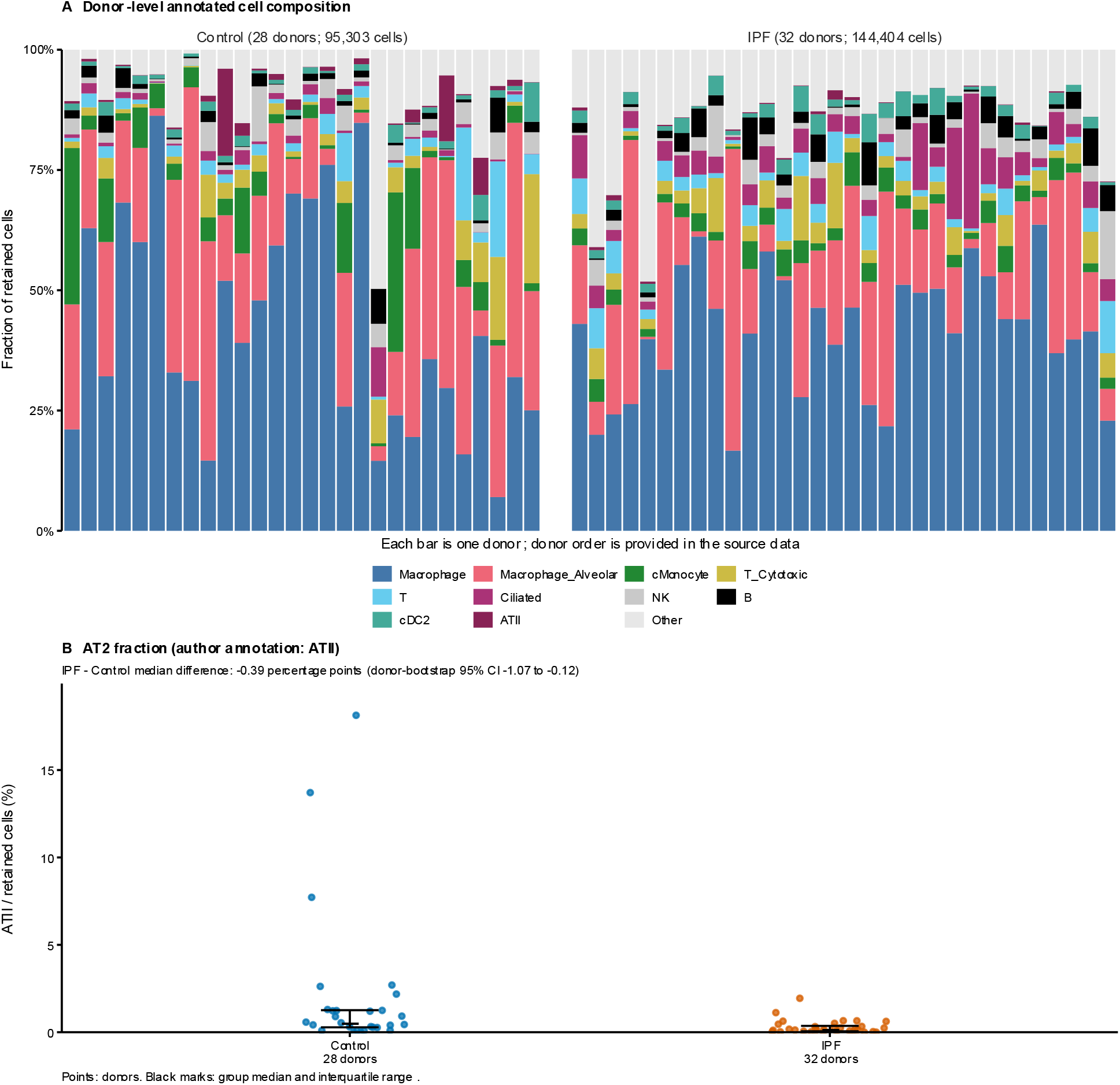
Donor-level lung cell composition in GSE136831. (A) Cell-type composition within each donor (IPF, n = 32; control, n = 28); The ten labels with the highest pooled cell counts are shown separately; the remaining labels are grouped as Other. (B) The fraction of retained annotated cells assigned to alveolar type II (AT2) cells for each donor. Summary marks indicate the median and interquartile range; group sizes count donors rather than cells. These fractions describe captured and annotated cells, not absolute abundance in lung tissue.

## 6 Discussion

SFL turns a scientific figure example into a reusable asset with an explicit relationship to code, context, and provenance. The central design choice is to maintain that relationship across discovery, publication, and transfer into a project. Visual inspection remains the entry point, while identifiable releases and recorded materialization carry the selected example into a code-based workflow. The accompanying module collection demonstrates how the same organization can support an openly distributed plotting resource.

The comparison identifies complementary objects of interaction: generated charts and data transformations, operations that users inspect or edit, and reusable assets that persist across projects. LIDA, Data Formulator 2, WaitGPT, and DynaVis develop interfaces around chart creation, iteration, or operational control [8,9,10,11]. SFL emphasizes the selected asset entering a project, preserving its identity and supporting materials. This comparison motivates evaluation questions specific to reuse: whether users recover the intended example, understand its inputs, retain provenance during adaptation, and transfer it into their own environment.

The division between client and resource collection also supports incremental development. New examples can be added to the open collection without embedding a new analysis engine in the client. Researchers can maintain local assets that reflect their own workflows while still drawing on shared modules. This makes the image–code relationship, rather than a particular host or model, the stable object around which reuse is organized.

The application case verifies the handoff from template selection to a checkable analysis output in one public lung cohort. It complements the software tests, which used synthetic protocol fixtures on Windows. The case did not test user efficiency, comparison with another tool, independent cohorts, or transfer across hosts. Its exploratory proportions depend on capture and annotation and are not evidence of an absolute change in tissue cell abundance or a disease mechanism.

## 7 Conclusion

Scientific Figure Library connects visual discovery with reusable plotting materials. Its local asset model preserves figure–code relationships, its provider interface brings independent collections into a common workflow, and its confirmation and materialization steps carry a selected example into a research project. Together with Open Figure Modules, SFL offers a modular foundation for collecting, sharing, and adapting scientific visualization knowledge across research environments.

## Supporting information

Supporting files: reproduction scripts, source data, and software evidence

## Code and resource availability

The SFL client is available at https://github.com/xuzhougeng/ScientificFigureLibrary [14]. The Open Figure Modules resource is available at https://github.com/jarxunlai/ScientificFigureLibrary-personal [15]. The examined source snapshots, resource-inventory procedure, and software-check records are documented in Supplementary Material S1 and its accompanying evidence files. The manuscript SVG schematics are included with the supporting files. The original IPF Cell Atlas metadata are available at GEO accession GSE136831 [23]. The review supplement includes derived donor-level counts, figure source data, reproducibility scripts, and the SFL materialization record for this case.

SFL client code is distributed under the MIT license. Resource licenses are specified per module; the three representative modules described here use MIT for code and CC BY 4.0 for synthetic content and documentation. The resource includes source modules, example inputs, metadata, previews, and downloadable archives [14,15].

## Author contributions

Jiaxuan Lai: Conceptualization, Methodology, Software, Data curation, Resources, Validation, Writing – original draft, and Writing – review & editing.

Shiming Xie assisted with writing the manuscript.

Yu Qiao, Xi Chen, Zhougeng Xu, and Shipeng Guo provided advice on the system design and architecture.

## Ethics and data use

This work used software, publicly available figure resources, synthetic examples, and secondary analysis of publicly deposited cell annotations from GSE136831 [23]. No new participants were recruited, no new specimens were collected, and no identifying information was used in this analysis. The original sample collection is described by Adams et al. [22].

## Acknowledgements

None.

## AI assistance

OpenAI Codex assisted with literature searches, source-code review, manuscript drafting and translation, SVG schematics, and implementation and verification of the public-data case study. The authors are responsible for the final content.

