## Supplementary Material S1 for "Scientific Figure Library connects visual examples with reusable plotting code"

### Source snapshots and supporting software records

This supplement documents the source inspection, resource inventory, and software checks reported in Scientific Figure Library connects visual examples with reusable plotting code. It accompanies the manuscript with the evidence needed to identify the inspected materials and understand the scope of the checks.

### S1 Source snapshots

The SFL client was examined at source commit `7257d4b1e488b686187e467579beb7d2686d52cc`. The repository contains the storage implementation, shared library service, provider interfaces, protocol documentation, local-client architecture, and automated tests. The inspected source is accessible through the fixed SFL source snapshot.

Open Figure Modules was inspected at source commit `bb9aa9c75bc92c9e43ed941823dfa3c001bedcd1`. The resource is accessible through the fixed Open Figure Modules snapshot. Repository and manifest inspection were performed on 5 November 2026. These identifiers locate the materials supporting this manuscript; they are not claims about subsequent repository changes.

### S2 Resource inventory

The complete repository tree response was inspected and was not truncated. Direct module directories under `modules/` were compared by identifier with the entries in `catalog/archive-manifest.json`. There were 182 module directories, 182 manifest entries with unique module identifiers, and 182 corresponding ZIP archives. The directory and manifest identifier sets matched. Each module directory included a primary preview, a browsing thumbnail, and a module manifest. These counts describe resource objects, not independent biological datasets or validated analyses.

The machine-readable inventory summary is supplied as `resource_inventory.json`. The module identifier list and public module URLs are supplied as `module_index.json`. The repository README contained an older prose count; the manuscript uses the directly inspected directory and archive-manifest inventory.

Three representative modules were examined through their descriptions, manifests, input schemas, actual CSV inputs, provenance records, and organized R scripts. Plotting scripts were not rerun during this resource inspection.

The sample-cell-type composition module, `sc-celltype-sample-stacked-proportion`, contained a 63-row, four-column synthetic count table with sample, group, cell type, and count fields. Its script calculates sample-level proportions and totals and constructs grouped composition displays.

The `neighbourhood-abundance-beeswarm` module contained two synthetic tables with 7,400 and 3,780 rows, respectively, each with four columns. They provide precomputed neighbourhood effects for plotting. The inspected entry point visualizes these supplied values rather than running the upstream neighbourhood differential-abundance analysis.

The `volcano-inset-enrichment-bars` module contained a 2,600-row, three-column gene-level table and a 12-row, four-column functional-summary table. The code constructs coordinated displays from supplied or illustrative synthetic values; it does not carry out the upstream differential-expression or enrichment analysis. These three modules declare MIT code licensing and CC BY 4.0 for the identified synthetic content and documentation.

### S3 Automated software checks

The inspected client snapshot was checked in an isolated source clone on Windows 10.0.26200, with Node.js 24.16.0 and npm 11.13.0. The commands were `npm ci`, `npm test`, and `npm run test:smoke`. All three completed successfully. The retained test runner reported 483 tests: 480 passed, zero failed, and three were skipped.

The skipped cases concerned the Linux no-node launcher, POSIX literal argument quoting, and symlink preview-cache behavior. The build-and-smoke command included TypeScript checks, Web and backend builds, and the MCP smoke script. Smoke output reported 62 standard tools and exercised intake, immutable publication, exact-preview confirmation, materialization and checked replay, selected bundle operations, and diagnostics.

The test wrapper created isolated home and configuration directories. The smoke script created its own temporary Library and inputs. Neither path was directed at a live user Library. The smoke fixture used one-pixel PNGs, a small R-code string, and simulated execution-evidence records. It did not invoke R to generate a scientific plot. Accordingly, the retained outcomes describe software behavior under the exercised fixtures.

To repeat the checks, obtain the identified source snapshot, install the recorded Node.js and npm environment, and run the same three commands from the source root. Keep the repository's isolation wrapper and lockfile. The accompanying `logs/` files preserve the original command outputs and were checked for local user/workspace prefixes and credential-like strings before packaging. The results were not rerun or reclassified when preparing this supplement.

### S4 Supporting file organization

The accompanying evidence package contains this supplement, `resource_inventory.json`, `module_index.json`, `verification_environment.json`, three command logs, and the three English SVG schematics used in the manuscript. The schematics explain architecture, confirmed reuse, and resource organization; they are not empirical plots. Public source and resource content remain accessible through the repositories above.

### S5 Public-data application and reproduction

We used the author-deposited cell annotation table and public sample metadata from GEO GSE136831. The IPF and control groups contributed 32 and 28 donors, respectively. The AT2 source label was `ATII`. The source counts include the author-labeled Multiplet records. The primary denominator excludes them (2,765 in IPF and 1,000 in controls); a sensitivity analysis retains them. No sequencing reads, expression matrix, reclustering, or individual clinical covariates were used. The case bundle supplies acquisition and processing scripts, derived count tables, figure source data, display-category mapping, and the audit of donor and denominator mappings.

The template and software versions used for this particular case are recorded with its materialization receipt. The evidence identifies the selected template and the project-specific adaptation separately. Run the case steps in the order specified in the accompanying README; the figure code runs in R outside SFL. The descriptive bootstrap resampled donors separately within groups (10,000 replicates; seed 136831), using the 2.5th and 97.5th percentiles. The interval does not adjust for clinical covariates or sampling bias.
