## Supporting files: reproduction scripts, source data, and software evidence for "Scientific Figure Library connects visual examples with reusable plotting code": 03_at2_fraction.pdf

**B AT2 fraction (author annotation: ATII)**

IPF - Control median difference: -0.39 percentage points (donor-bootstrap 95% CI -1.07 to -0.12)

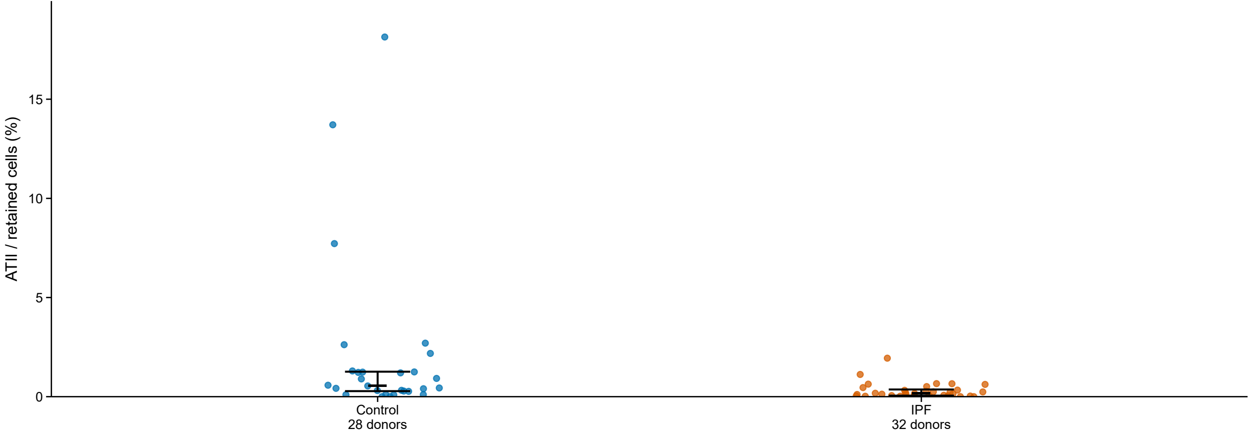


Points: donors. Black marks: group median and interquartile range.
