## Supporting files: reproduction scripts, source data, and software evidence for "Scientific Figure Library connects visual examples with reusable plotting code": 03_donor_composition.pdf

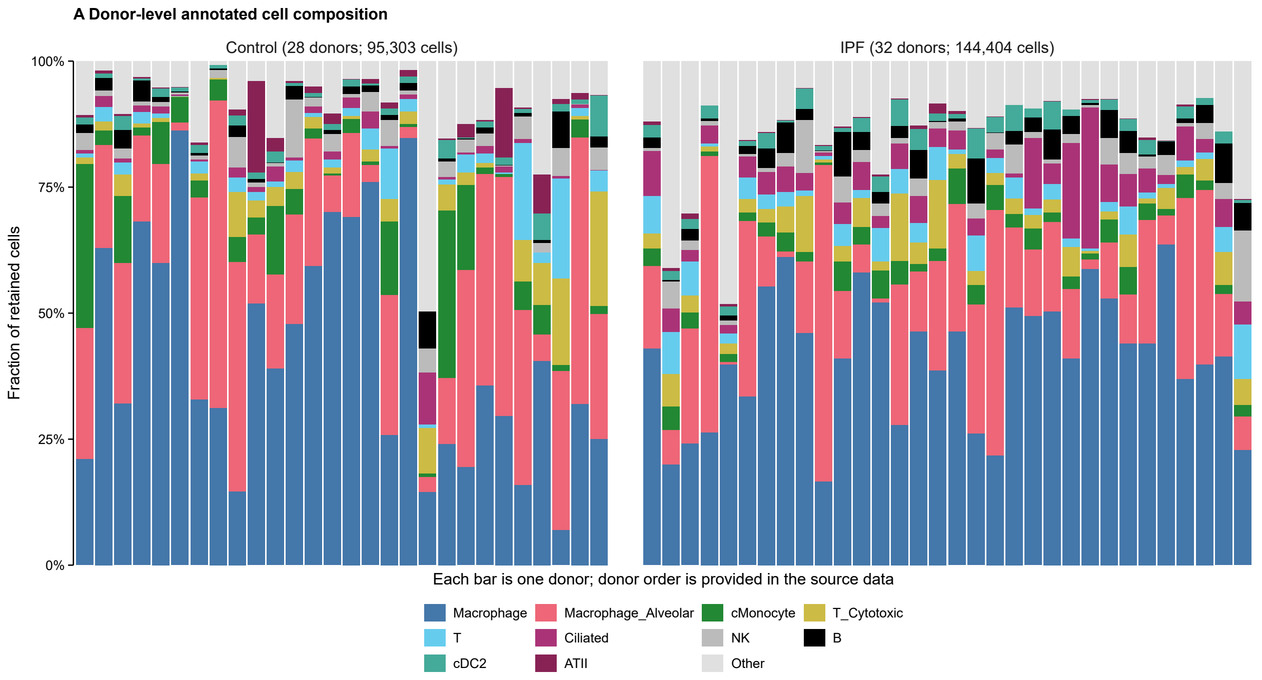
A Donor-level annotated cell composition
Control (28 donors; 95,303 cells)
IPF (32 donors; 144,404 cells)
Fraction of retained cells
100%
75%
50%
25%
0%
Each bar is one donor; donor order is provided in the source data
Macrophage
Macrophage_Alveolar
cMonocyte
T_Cytotoxic
T
Ciliated
NK
B
cDC2
ATII
Other
