## Supporting files: reproduction scripts, source data, and software evidence for "Scientific Figure Library connects visual examples with reusable plotting code": fig4_ipf_composition.pdf

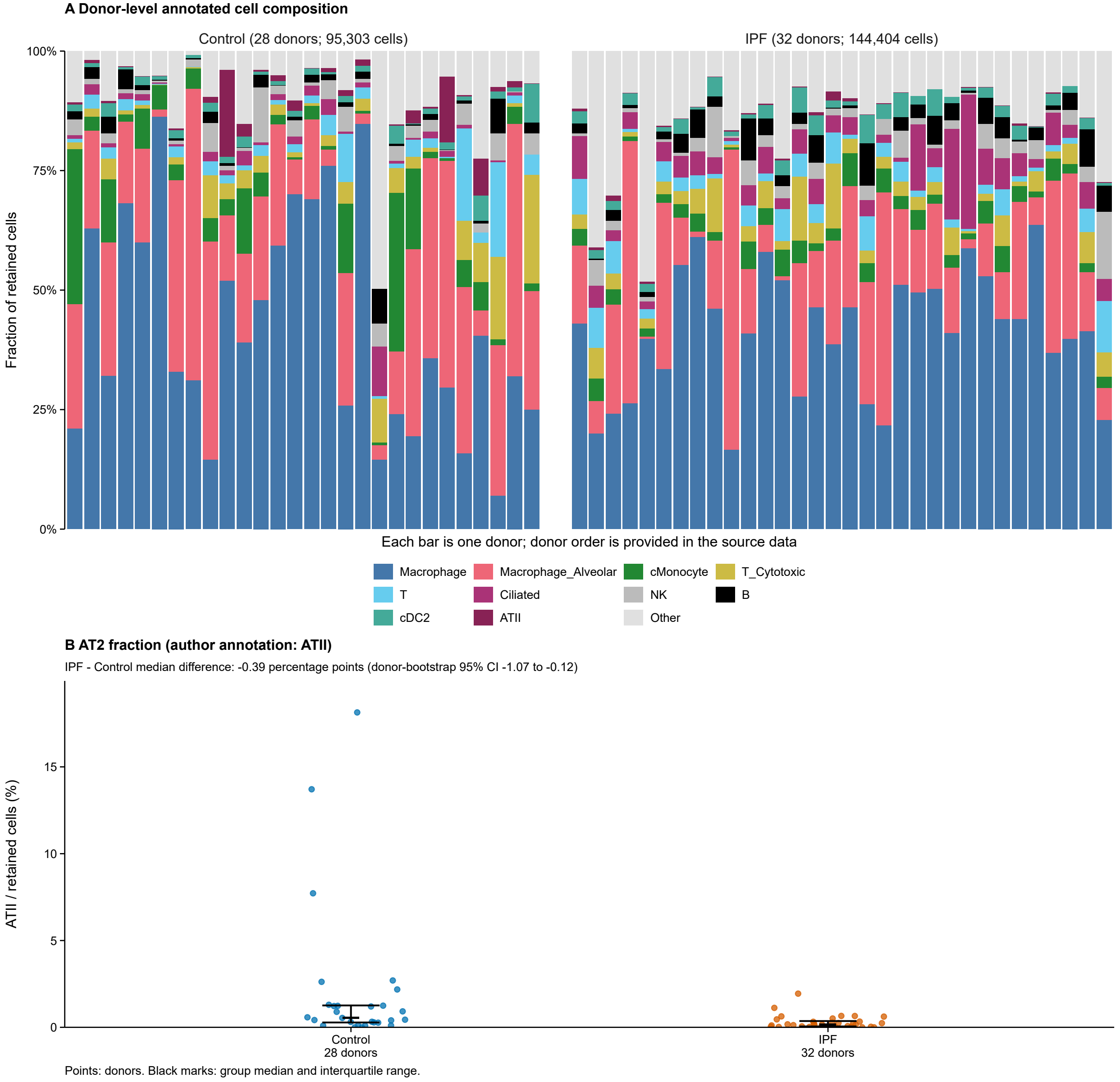
A Donor-level annotated cell composition
Control (28 donors; 95,303 cells)
IPF (32 donors; 144,404 cells)
100%
75%
50%
25%
0%
Fraction of retained cells
Each bar is one donor; donor order is provided in the source data
Macrophage
Macrophage_Alveolar
cMonocyte
T_Cytotoxic
T
Ciliated
NK
B
cDC2
ATII
Other
B AT2 fraction (author annotation: ATII)
IPF - Control median difference: -0.39 percentage points (donor-bootstrap 95% CI -1.07 to -0.12)
ATII / retained cells (%)
15
10
5
0
Control
28 donors
IPF
32 donors
Points: donors. Black marks: group median and interquartile range.
