## Supplementary figures and images for "Scientific Figure Library connects visual examples with reusable plotting code"

### 03_at2_fraction.png

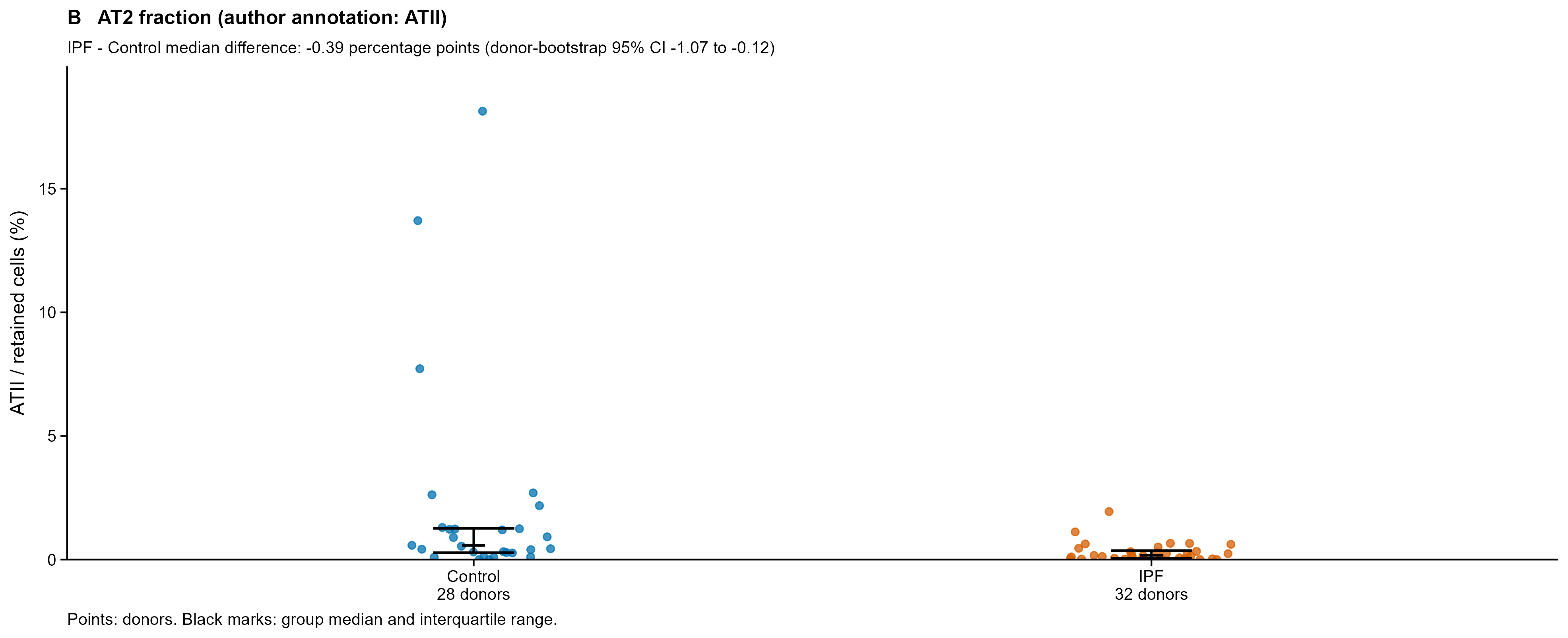

### 03_donor_composition.png

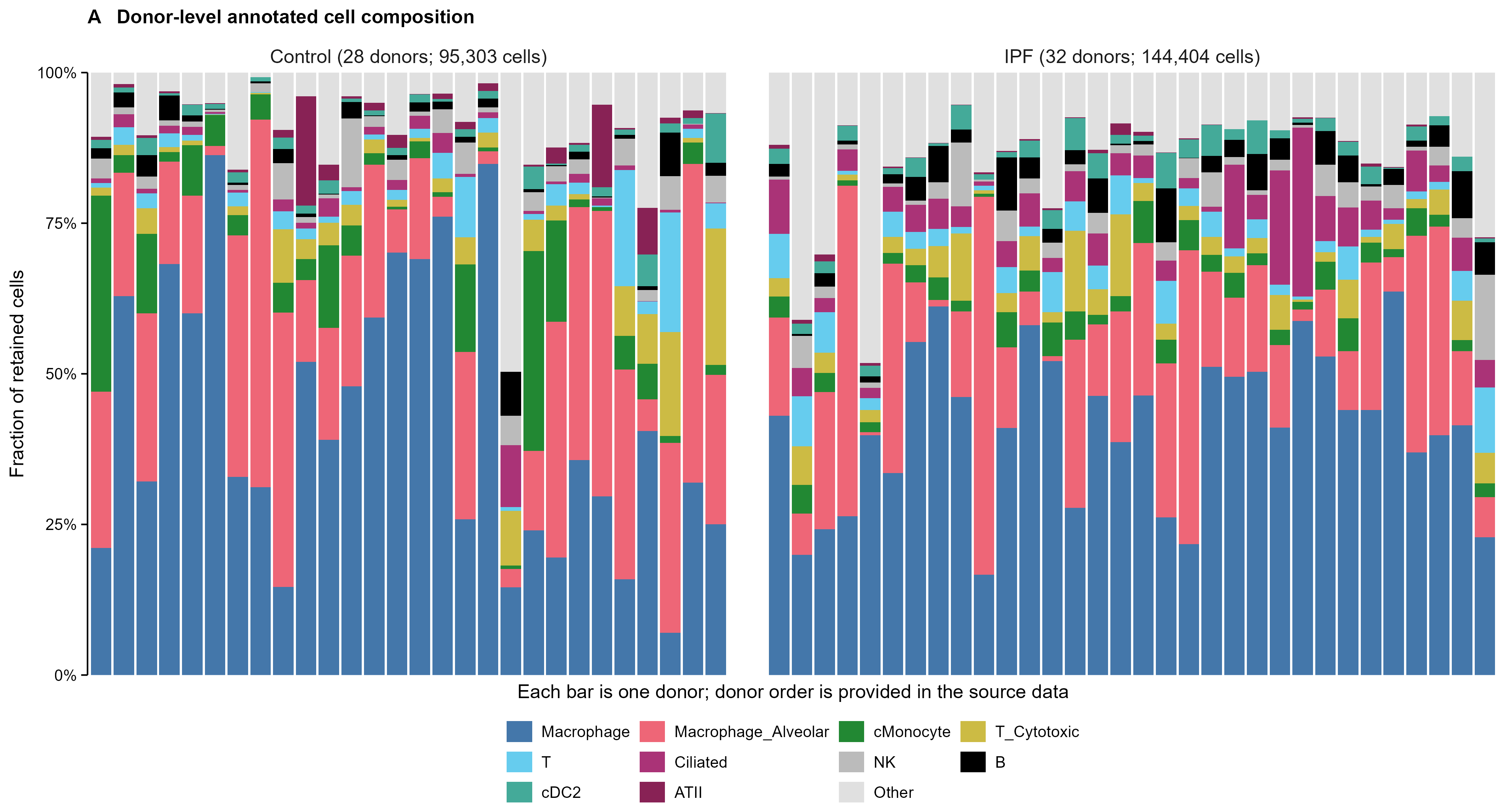

### fig4_ipf_composition.png

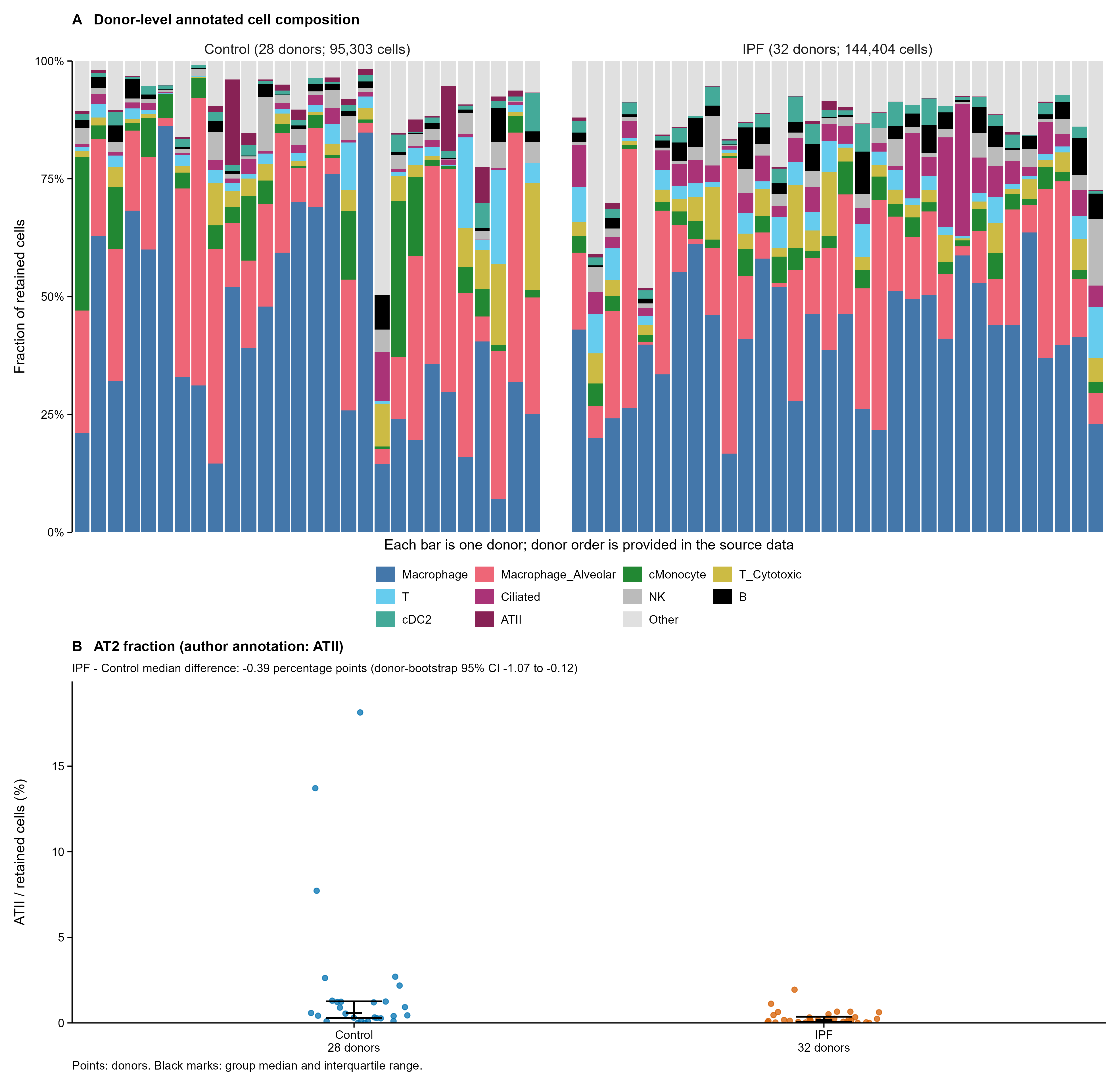
